# Metabolites alleviate staphylococcal bloodstream infection in a NO-dependent manner via arginase inhibition

**DOI:** 10.1101/2020.03.02.974345

**Authors:** Rui Pang, Yu-bin Su, Hua Zhou, Xinhai Chen

## Abstract

*Staphylococcus aureus* is a notorious bacterial pathogen that often causes soft tissue and bloodstream infections and invariably garners resistance mechanisms against new antibiotics. Host innate immunity modulated by metabolites has been proved as a powerful strategy against bacterial infections. However, few studies focus on the application of this strategy against *S. aureus* infection. Here, we identified four metabolite biomarkers, L-proline, L-isoleucine, L-leucine, and L-valine (PILV), by a metabolomics study. In animal models of *S. aureus* bloodstream infection, exogenous administration of each metabolite or PILV shows an anti-infective effect, while PILV treatment has higher protection than individual metabolite treatment. Each metabolite targets nitric oxide (NO) to kill *S. aureus* via arginase inhibition, and PILV exhibits additive inhibition of arginase activity that causes further killing. This suppression also contributes to the metabolite-mediated phagocytic killing of *S. aureus* in human blood. Our finding demonstrates the metabolite-mediated innate immunity as a therapeutic intervention for *S. aureus* infection.

---

The pathogen *Staphylococcus aureus* is both a human commensal and a significant cause of hospital- and community-acquired diseases including soft tissue infections, pneumonia, osteomyelitis, septic arthritis, bacteremia, endocarditis, and sepsis (1–3). The asymptomatic colonization is common; however, 80% invasive *S. aureus* strains isolated from the blood of patients with *S. aureus* bacteremia are genetically indistinguishable to the nasal strains detected at admission (4). Because of the high incidence of hospital-acquired infection, antibiotics are employed both for *S. aureus* decolonization and prophylaxis of nosocomial disease (5, 6). However, the emergence and spread of drug-resistant strains, designated MRSA (methicillin-resistant *S. aureus*), led to increased therapeutic failure and mortality rates due to *S. aureus* infections (6). Therefore, new approaches are especially needed for treating such infections in the clinic. One possible approach would be to enhance the innate immune response of the infected host, restoring the defense ability to kill the bacterial pathogen in a relatively risk-free manner (7).

Similar to other bacterial infections (8, 9), *S. aureus* infection causes several metabolisms changed pronouncedly in the host, which contain oxidative phosphorylation, aerobic glycolysis, and amino acid and fatty acid metabolisms (10–14). A growing body of recent studies shows that bacterial infection-induced shift of metabolisms has two leading roles, which either facilitate the bacterial invasion or benefits to the immune responses against bacterial infection. Host central carbon metabolism is capable of being strongly disturbed by *S. aureus*, which activates autophagy by increasing the phosphorylation of AMP-activated protein kinase and extracellular signal-regulated kinase, thereby meeting the staphylococcal invasion (15). Furthermore, the internalization of *S. aureus* employs two own pathways to destroy the host arginine metabolism, limiting the production of nitric oxide (NO), which serves in the host’s antibacterial defense, and eventually inducing death of the host cell (16, 17). On the other hand, studies focusing on the cross-talking between metabolic regulation and immune system reveal an active role of metabolic regulation in controlling the pathogenic bacteria. In several bacterial infection models, the host that survive the infection display distinctive metabolic pathways (18–23). Numerous metabolites identified from these metabolic pathways related to the survivors can be immunoregulators that modulate the function of the immune system via various mechanisms, including the activation of PI3K/Akt1, elevated expression of cytokines, and promotion of NO production (18–23). However, few studies are operated to investigate whether the modulation of host innate immunity by metabolites is a valuable strategy against staphylococcal infection.

Here, we used a gas chromatography-mass spectrometry (GC-MS) to identify metabolites from BALB/c mice infected by three increasing sublethal doses of *S. aureus* strain Newman. The results suggest that four metabolites (L-proline, L-isoleucine, L-leucine, and L-valine) target the NO production to kill *S. aureus*, which may aid in the development of therapeutic interventions that can improve the outcome of MRSA infection.

## Results

### GC-MS-based metabolomics identifies host metabolites relating to *S. aureus* infection

To exploit anti-infection metabolites from the host, metabolic profiling with different degrees of anti-infection should be established. We hypothesized that different sublethal infection doses would induce different degrees of anti-infection in the host. Therefore, BALB/c mice were intravenously challenged with Low, moderate, or high sublethal dose of *S. aureus* Newman or with PBS. 12 h later, plasma samples were separated from these challenged mice, and the GC-MS-based approach was used to identify crucial metabolites. A total of 72 metabolites were detected in each sample and displayed as a heat map (Fig. 1A). The heat map showed that the majority of metabolites were changed in abundance, suggesting that *S. aureus* infection altered the mouse plasma metabolome. No infection, low dose, moderate dose, and high dose groups could be distinguished by principal component analysis (PCA) using 72 metabolites (Fig. 1B), which demonstrated our hypothesis that hosts infected by different sublethal doses drive different metabolic profiling of anti-infection. After assaying, 48, 44, and 27 differential metabolites were respectively detected by comparisons of no infection and low dose group, of low and moderate dose groups, and of moderate and high dose groups (Fig. 1C), among which 14 metabolites were shared (Fig. 1C and 1D). A subset of six metabolites, including L-leucine, L-proline, L-isoleucine, monolinolein, L-valine, and eicosanoic acid was significantly increased on infection from low dose to moderate dose to high dose (Fig. 1D). These metabolites could serve as potential anti-infection biomarkers for *S. aureus* infection. Additionally, 14 shared metabolites enriched for four pathways containing aminoacyl-tRNA biosynthesis, citrate cycle, valine, leucine, and isoleucine degradation, and valine, leucine, and isoleucine biosynthesis (*P* < 0.05) (Fig. 1E). Out of six metabolite biomarkers, four metabolites including L-leucine, L-proline, L-isoleucine, and L-valine were enriched in aminoacyl-tRNA biosynthesis, valine, leucine, and isoleucine degradation, and valine, leucine, and isoleucine biosynthesis, thereby motivating us to investigate these metabolite biomarkers in greater detail (Fig. 1F).

**Figure 1.**
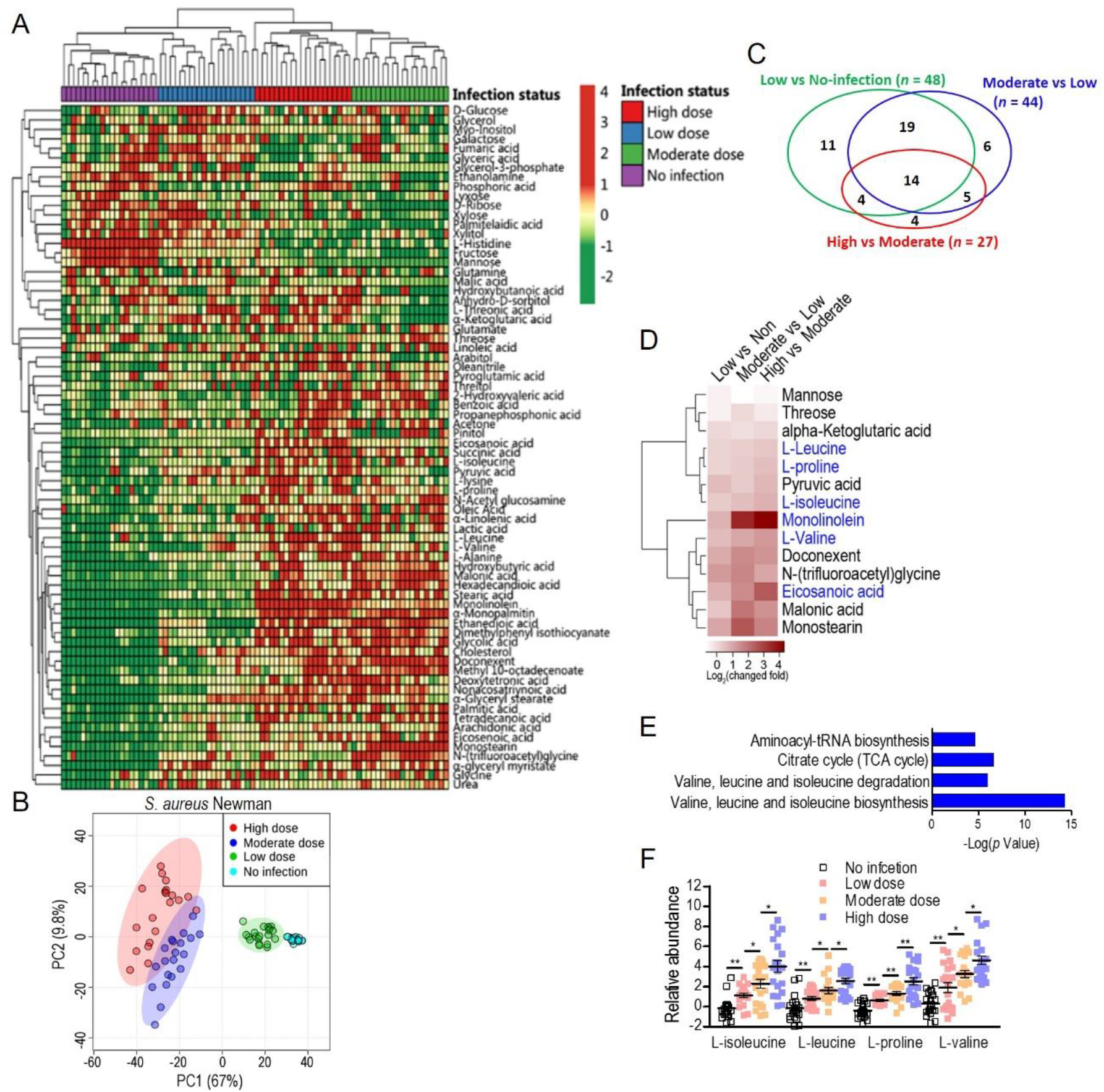
Serum metabolome analysis by GC-MS reveals L-isoleucine, L-leucine, L-proline, and L-valine as potential anti-infection metabolites against *S. aureus* infection. **(A)** The heat map showed the relative abundance of total 72 metabolites in serum samples from mice infected by low dose (0.3 × 10^7^ CFU), moderate dose (0.7 × 10^7^ CFU), and high dose (1 × 10^7^ CFU) of *S. aureus* Newman strain, respectively, or PBS as no-infection control. **(B)** Principal component analysis (PCA) led to the metabolomics discrimination among no-infection control, low dose, moderate dose, and high dose groups. **(C)** Venn diagram of 48 differential metabolites from the comparison of the low-dose group (Low) to the no-infection group, 44 differential metabolites from the comparison of the moderate-dose group (Moderate) to Low, and 27 differential metabolites from the comparison of the high-dose group (High) to Moderate. **(D)** Heat map representation of unsupervised hierarchical clustering of fourteen overlapped metabolites in **(C)**. **(E)** Pathway enrichment analysis of fourteen overlapped metabolites. A horizontal histogram was selected to show the impact of the enriched pathway with *P* values < 0.01. **(F)** The abundance of L-isoleucine, L-leucine, L-proline, and L-valine in no-infection, Low, Moderate, and High dose groups. Error bars ± SEM, * *P* < 0.05 and ** *P* < 0.01.

### Exogenous metabolites show the anti-infective effect on *S. aureus* infection

To examine the potential anti-infective role of L-leucine, L-proline, L-isoleucine, or L-valine in *vivo*, cohorts of mice were intravenously infected with *S. aureus* Newman [1 × 10^7^ colony-forming units (CFU)] and injected with each metabolite (0.5 g kg^−1^) or sterile saline (no metabolite control) daily. Compared to mice administrated with sterile saline, metabolite-injected mice significantly declined the bodyweight loss during *S. aureus* infection (Fig. 2A). When renal tissues were analyzed for bacterial burdens and histopathology and compared with saline-administrated animals, mice given metabolite displayed markedly reduced staphylococcal loads and numbers of abscess lesions (Fig. 2B and 2C). Each metabolite vaccination also provided distinct protection against lethal bloodstream infection with USA300 (2 × 10^8^ CFU) and MRSA252 (2 × 10^9^ CFU) (Fig. 2D and 2E). Besides, there was no difference in the recovery of body weight, bacterial loads, abscess numbers, and survival among each metabolite administration (Fig. 2A-2E). More importantly, we surprisingly found that combined administration of L-proline, L-isoleucine, L-leucine and L-valine (PILV) is capable of promoting survival further (Fig. 2D and 2E), which were not found by administration of 4-fold higher concentration of L-proline (2.0 g kg^−1^) (Fig. S1), indicating the characteristically synergetic effect of four metabolites against staphylococcal infection. Thus, based on these data, we find that L-leucine, L-proline, L-isoleucine, or L-valine has an anti-infective function during *S. aureus* infection and PILV combination treatment would further improve the therapeutic effect on MRSA infection.

**Figure. 2.**
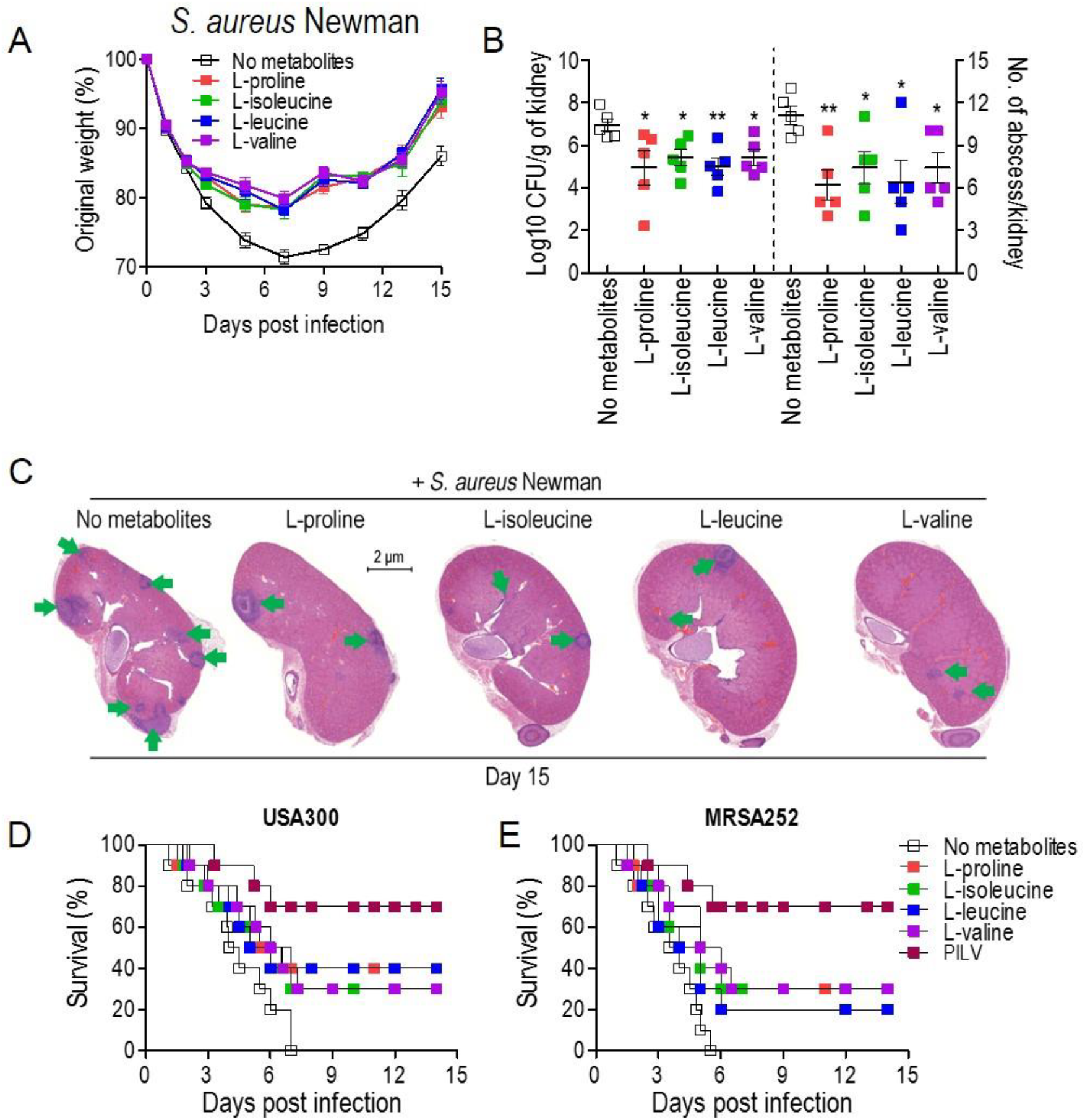
The administration of single metabolite or metabolite combination (named PILV) protects mice against *S. aureus* bloodstream infection. **(A to C)** The treatment of each metabolite, L-proline, L-isoleucine, L-leucine, or L-valine, rescued body weight loss **(A)** and reduced renal bacterial loads and abscess numbers **(B and C)** from mice infected by *S. aureus* Newman strain. Weight was recorded daily and reported as % of initial weight. Fifteen days post infection, kidneys (*n* = 5) were removed, and either ground for enumeration of CFU/g tissue or fixed for counting of surface abscesses **(B)**. Fixed kidneys were additionally thin sectioned and then stained with hematoxylin and eosin (H&E) for internal abscesses **(C)**. Green arrows point to internal abscesses in the kidney. **(D and E)** The treatment of single metabolite or metabolite combination (L-proline, L-isoleucine, L-leucine, and L-valine, designated as PILV) protected mice (BALB/c, *n* = 20) against lethal bloodstream infection with *S. aureus* USA300 **(D)** and MRSA252 **(E)**. Survival was monitored over fourteen days. Data are represented as Error bars ± SEM. * *P* < 0.05 and ** *P* < 0.01.

### Metabolites boost NO production by inhibition of arginase *in vivo*

Lipopolysaccharide (LPS), a component of the cell wall of Gram-negative bacteria, simultaneously induces expression of arginase (Arg) and NO synthase (NOS) in the host. However, it is unclear whether Gram-positive bacteria, *S. aureus*, is also able to induce both Arg and NOS expressions simultaneously. Thus we first measured NO production in serum samples and expression level of two Arg (cytoplasmic and mitochondrial arginase, designated as Arg1 and Arg2, respectively) and three NOS isozymes (neuronal, inducible, and endothelial NOS, designated as NOS1, 2, and 3, respectively) in tissues and blood upon *S. aureus* infection. Three days post sublethal infection of *S. aureus* Newman, NO production was enhanced in a dose-dependent manner (Fig. 3A). Furthermore, intravenous infection of *S. aureus* USA300 triggered the expression of all arginase and NOS isozymes and increased the NO production and arginase activity in mouse tissues (liver and kidney) and blood (or serum) except unchanged NOS3 expression in the blood (Fig. 3B to 3J). More interestingly, *S. aureus* infection induced more expression of Arg isozymes than NOS isozymes in tissues (Fig. 3B and 3C), suggesting that both Arg isozymes are predominant regulators of L-arginine since Arg and NOS compete with one another for L-arginine as an enzyme substrate. Then we asked whether L-leucine, L-proline, L-isoleucine, L-valine, or PILV have a mechanism that boosts NO production by blocking the arginase activity under the condition of *S. aureus* infection. Cohorts of mice were daily intraperitoneal injection of each metabolite with 0.5 g kg^−1^ or 2.0 g kg^−1^ or of PILV (0.5 g kg^−1^ for each metabolite) and infected by *S. aureus* after 6 hours post the first injection of metabolites. On day 3, animals were euthanized, and their blood and tissues (kidney and liver) were collected for measurements of NO production, arginase activity, and urea level. Mice that had received PILV held the highest level of NO in serum and tissue and the lowest activity of arginase and level of urea in serum, followed by those that had received one metabolite or 4-fold higher concentration of that metabolite (Fig. 3E to 3K). In the absence of *S. aureus* infection, 0.5 g kg^−1^ or 2.0 g kg^−1^ of each metabolite or PILV (0.5 g kg^−1^ for each metabolite) showed no impact on NO production (Fig. S2). These data suggest that L-leucine, L-proline, L-isoleucine, or L-valine can strengthen NO production, and PILV combination therapy has an additive effect on NO production through further inhibition of arginase.

**Figure 3.**
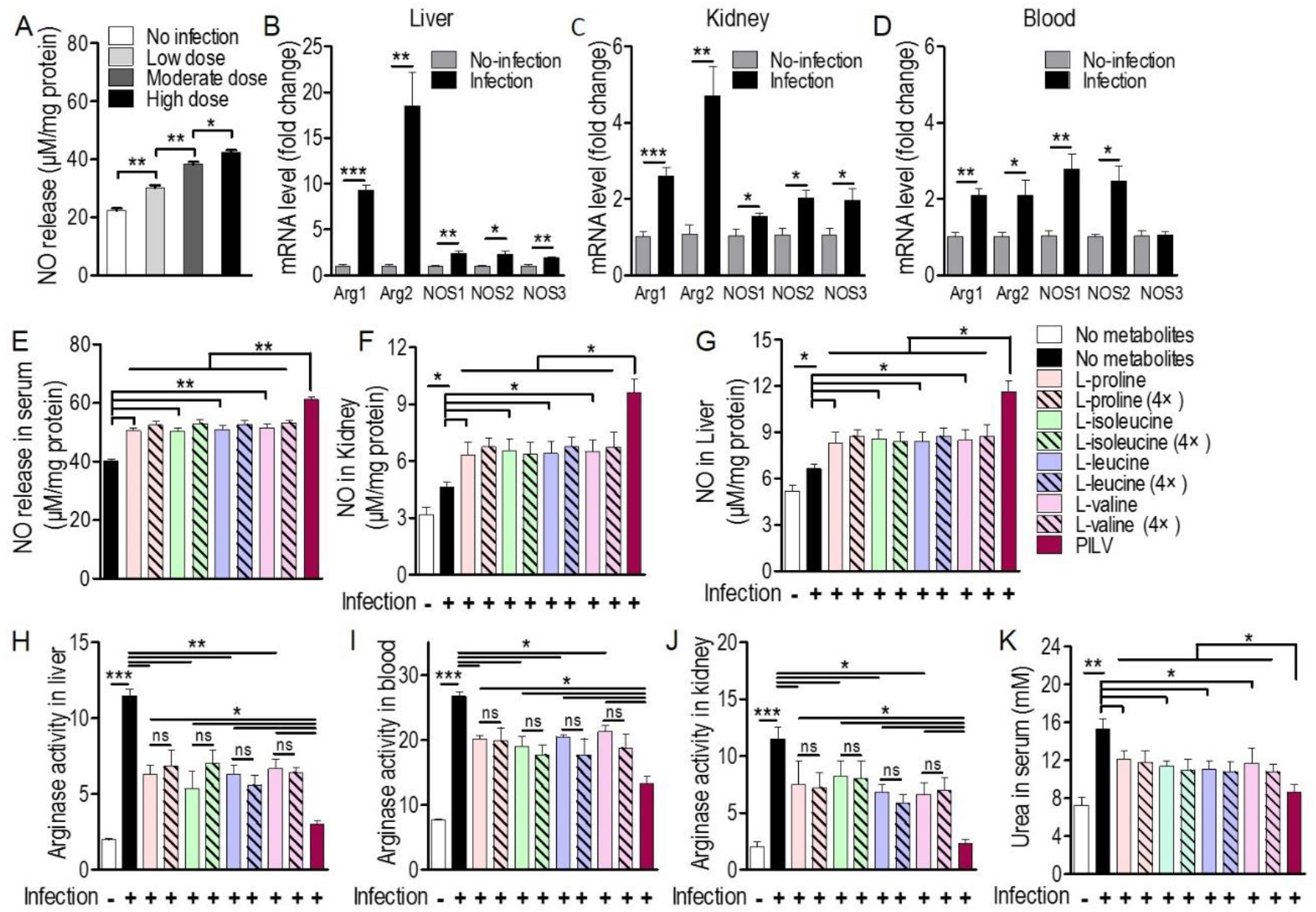
Metabolites promote NO production by inhibition of arginase in mice. **(A)** Staphylococcal infection enhanced NO release in a dose-dependent manner. Three sublethal doses from low to high (0.3 × 10^7^, 0.7 × 10^7^, and 1 × 10^7^ CFU/mouse) of *S. aureus* Newman strain was used to intravenously infect mice (*n* = 5 for each dose group). Three days post infection, mouse serum was collected and subjected to the measurement of NO release. **(B to D)** The mRNA levels of two arginase (Arg) and three NO synthase (NOS) isoforms in mouse liver, kidney, and blood were up-regulated by a sublethal staphylococcal infection. *S. aureus* USA300 (5 × 10^6^ CFU/mouse) was used to infect BALB/c mice (*n* = 5). Three days post infection, mouse liver, kidney, and blood were collected and then subjected to estimate the transcriptional expression of Agr and NOS. Agr1 and Agr2 are cytoplasmic and mitochondrial arginase, respectively. NOS1, NOS2, and NOS3 are neuronal, inducible, and endothelial NOS, respectively. **(E to J)** Upon staphylococcal infection, single metabolite or PILV treatment increased NO production by inhibiting arginase activity in mouse blood and tissues. NO release was increased by single metabolite treatment and further boosted by PILV treatment in mouse serum **(E)**, kidney **(F)**, and liver **(G)**. Meanwhile, arginase activity was decreased by single metabolite treatment and further declined by PILV treatment in the liver **(H)**, blood **(I)**, and kidney **(J)**. **(K)** Staphylococcal infection-induced urea content was reduced by single metabolite treatment and further decreased by PILV treatment. Data are represented as Error bars ± SEM. * *P* < 0.05 and ** *P* < 0.01.

### Metabolites-induced NO protects mice against *S. aureus* infection

Because of the positive correlation between higher protection and NO production of PILV treatment, we presumed that PILV-induced NO production would be responsible for higher protection. To test this, we used one competitive arginase inhibitor called BEC, which shows no effect on NO production and urea level under physiological condition while enhancing NO production and decreasing urea level in serum samples of *S. aureus*-infected mice (Fig. 4A and 4B). Further survival assay presented that BEC protected against lethal challenge with MRSA strain USA300 (Fig. 4C). These data indicate that increasing NO production has the benefit of getting rid of MRSA infection. Then we investigated the effects of two NO inhibitors, l-NMMA, and l-NAME, on PILV-induced NO production and survival. As expected, the inhibitors significantly suppressed NO production induced by *S. aureus* infection in the absence or presence of PILV (Fig. 4D). The mouse survival caused by *S. aureus* infection or enhanced by PILV administration was all reduced by the NO inhibitors (Fig. 4E). Together, these data prove that PILV-induced NO production confers protection against staphylococcal disease.

**Figure 4.**
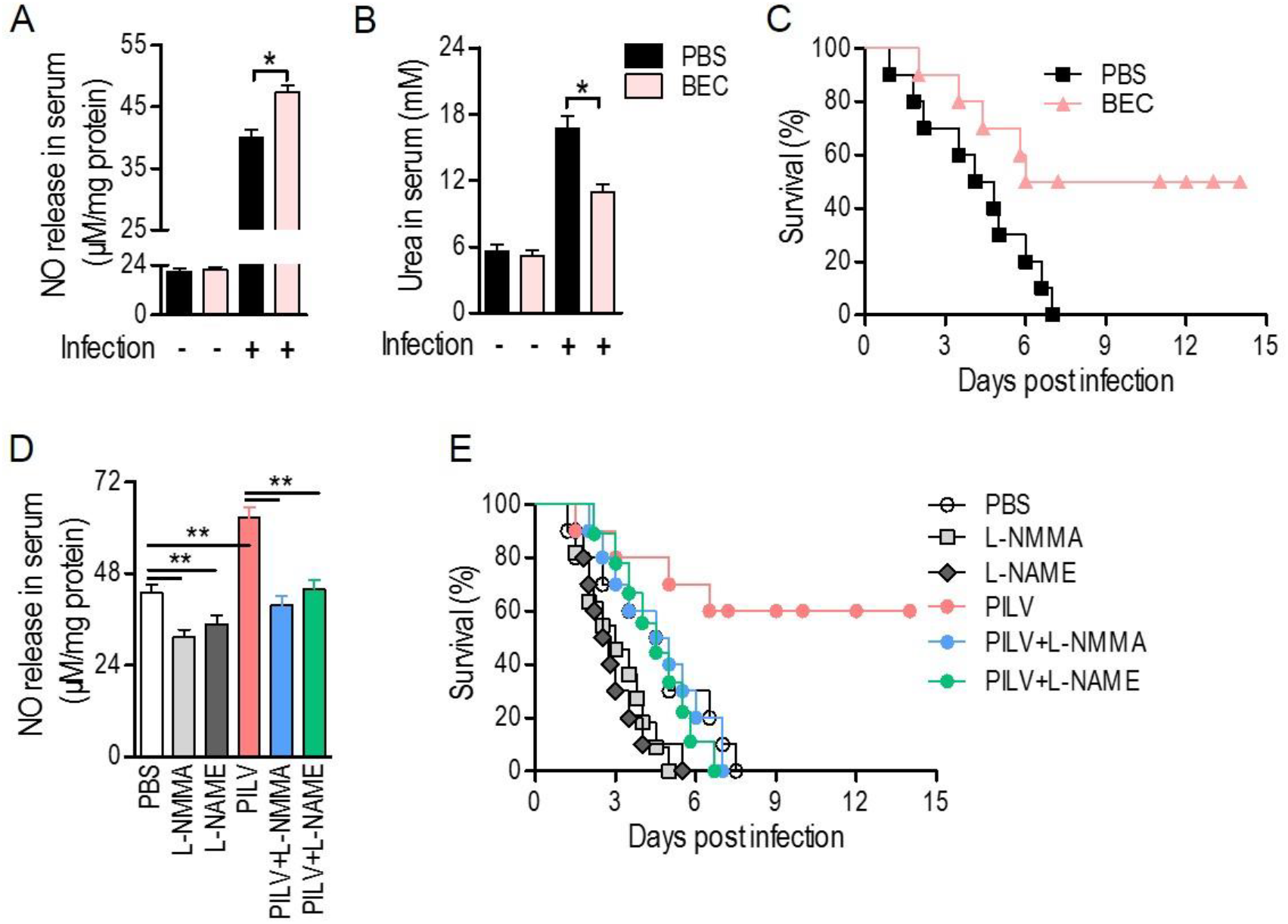
Metabolites-induced NO confers protection against staphylococcal infection. **(A to C)** A competitive arginase inhibitor, BEC (S-(2-boronoethyl)-L-cysteine), was able to induce NO release **(A)**, inhibit urea production **(B)**, and protect mice against lethal bloodstream infection with *S. aureus* USA300 **(C)**. **(D and E)** Both NO inhibitors, L-NMMA and L-NAME, all blocked PILV-induced NO release **(D)** and removed PILV-induced protection **(E)**. Six hours after injection of BEC (50 mg kg^−1^), L-NMMA (40 mg kg^−1^), L-NAME (40 mg kg^−1^), PILV, PILV plus L-NMMA (40 mg kg^−1^), or PILV puls L-NAME (40 mg kg^−1^), BALB/c mice (*n* = 30) were lethally challenged by *S. aureus* USA300 and then divided into two subgroups. One subgroup (*n* = 10) was used for the measurement of NO and urea, and another (*n* = 20) was used for observation of survival. Five survival mice (*n* = 10) at three days post infection were euthanized to measure the NO and urea production in serum. Survival was monitored over fourteen days. Data are represented as Error bars ± SEM. * *P* < 0.05 and ** *P* < 0.01.

### Metabolites increase phagocytic killing of *S. aureus* in a NO-dependent manner

We next asked if the PILV has a function in human blood. *S. aureus* opsonophagocytic killing (OPK) was measured in human blood infected with 5 × 10^6^ CFU Newman for 60 min. Before that, blood was pretreated with heat-killed *S. aureus* Newman for 30 min at 37°C. When added to blood samples, PILV reduced the bacterial load to 75% (Fig. 5A), indicating the anti-infective role of PILV in human blood. Treatment of human blood with NO inhibitor abolished OPK of Newman in the absence or presence of PILV (Fig. 5A). Similar results were found when measuring the OPK of *S. aureus* in mouse blood (Fig. 5B). Further, the specific phagocytosis of *S. aureus* Newman was determined in macrophage cell lines, RAW264.7, and differentiated U937 cells. As anticipated, NO inhibitor or cytochalasin D completely removed PILV-enhanced phagocytosis of *S. aureus* in either human or mouse macrophages (**Fig.5C and 5D**). Altogether, these data demonstrate that PILV promotes the phagocytic killing of *S. aureus* in a NO-dependent manner.

**Figure 5.**
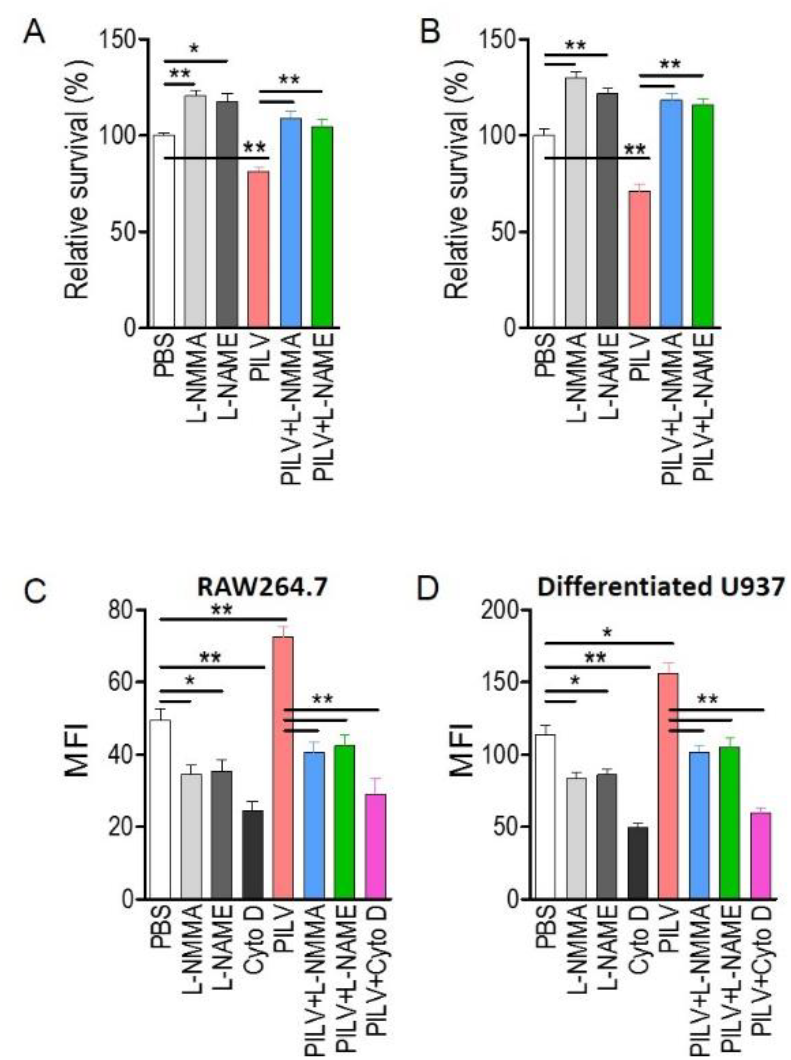
PILV enhances Opsonophagocytic killing (OPK) in a NO-dependent manner. **(A and B)** PILV promoted OPK of *S. aureus* in human **(A)** and mouse blood **(B)** through a NO-dependent manner. Anticoagulated and heat-killed *S.aureus* Newman-pretreated mouse and human blood was incubated with live S. aureus Newman (2.5 × 10^6^ CFU/ml blood for human blood assay, 2.5 × 10^5^ CFU/ml blood for mouse blood assay) in the presence of PBS, NO inhibitors, PILV, or PILV plus NO inhibitors for 60 min, and survival was measured (*n* = 5). **(C and D)** PILV increased the phagocytosis of FITC-conjugated *S. aureus* Newman in RAW 264.7 **(C)** and U937-derived macrophages **(D)**. RAW264.7 or U937-derived macrophages were pretreated with PBS, NO inhibitors, PILV, or PILV plus NO inhibitors in a serum-starved medium for 6 hours and then were co-incubated with FITC-conjugated *S. aureus* for an additional one hour. Bacterial uptake was measured by flow cytometry. Data are represented as Error bars ± SEM. * *P* < 0.05 and ** *P* < 0.01.

## Discussion

NO is a versatile effector that plays a central role both in the antimicrobial activity and immunomodulatory roles. During infection, the high cytotoxic NO level produced by innate immune cells of mammalian host limits pathogen growth (24). Although *S. aureus* has several genes for efficient NO detoxification, NO production is still critical for host resistance to staphylococcal disease (25–27). Besides the growth inhibition, NO can additionally target the Agr quorum sensing system to disrupt cell-to-cell communication of *S. aureus*, thereby suppressing the staphylococcal virulence (28). On the other hand, NO acts as a signaling messenger that promotes the growth and activity of immune cell types, including macrophages and neutrophils (24, 29). Inhibition of inducible NOS (iNOS) or total NOS in macrophages or peripheral blood neutrophils significantly blocks phagocytosis, intracellular killing, and increases the survival of *S. aureus* (23, 30–32). These observations result in the development of NO delivery systems that can harness the antimicrobial properties of this short-lived, evanescent gas (24, 33). Here we found another interesting way to enhance endogenous NO production in the presence of *S. aureus* infection through a combination therapy using four metabolites that were determined from a metabolomics study.

Metabolomics is an advanced technology that examines metabolic processes, identifies relevant biomarkers responsible for metabolic features, and discloses metabolic mechanisms. Analysis of the crucial metabolites in samples with different status has virtually become a significant part of improving the diagnosis, prognosis, and therapy of diseases (34). A platform for identifying metabolite biomarkers has been established in a mouse model of *Klebsiella pneumoniae* challenge (23). In that study, we garnered a potential anti-infection metabolite, L-valine, which presents an elevated level in surviving mice but a reduced level in dying mice. The supplementation of exogenous L-valine promotes the clearance of bacterial pathogens and enhances macrophage phagocytosis in a PI3K/Akt1 and NO-dependent manner. Using the same platform, we currently discovered four metabolite biomarkers from *S. aureus* bloodstream infection in an animal model. These metabolites are L-proline, L-isoleucine, L-leucine, and L-valine, the abundance of which becomes higher in plasma while infection dose increases, revealing the intriguing interrelationship between metabolites and infection status. The follow-up experiments not only evidence the contribution of metabolites in host resistance to staphylococcal infection but also provide the mechanistic link between metabolites and its anti-infection activity.

Using screening metabolites from differential metabolomics to combat bacterial infections have been well demonstrated in several studies. Plenty of metabolites, including glucose, malate, N-acetylglucosamine, Myo-inositol, linoleic acid, L-proline, L-valine, and L-leucine, show extended protection against bacterial infections (18–20, 22–24, 35, 36). However, the mechanistic investigations underlying their anti-infection properties are minimal. Earlier works showed the inhibitory effect of L-proline, L-isoleucine, L-leucine, and L-valine on arginase activities, and this inhibition is relatively specific as other amino acids, such as glycine, L-glutamine, do not influence arginase activities (37–39). Arginase inhibition by L-valine increases NO production in endothelial cells and macrophages in response to LPS treatment (23, 40). Specific elimination of arginase in macrophages favors host survival in *Toxoplasma gondii* infection and reduces the bacterial loads in the lung infection with *Mycobacterium tuberculosis* (41). Although *S. aureus* bloodstream infection co-induced expressions of NOS and arginase in the present study, the enzymatic role of arginase is distinctly more robust than that of NOS. Interestingly, L-proline, L-isoleucine, L-leucine, or L-valine is capable of boosting NO production via the inhibition of *S. aureus*-induced arginase activity in infected hosts. More importantly, the combined administration of L-proline, L-isoleucine, L-leucine, and L-valine has an additive effect on arginase inhibition, therefore providing the stronger protection against *S. aureus* infection largely through a mechanism that the more L-arginine is consumed by NOS to produce NO. There are two described isoforms of arginase, arginase I and arginase II, which are located on cytosol and mitochondria, respectively. Branched-chain amino acids (L-isoleucine, L-leucine, and L-valine) cause significant inhibition of cytosolic arginase I and only minor effect on mitochondrial arginase II, while L-proline has much more inhibition of mitochondrial arginase II than the cytosolic arginase II (37). This evidence probably explains why the more substantial effect of arginase inhibition only happens in combined administration of L-proline and branched-chain amino acids but not in the administration of 4-fold higher concentration of each that metabolite. Further investigation is required to determine whether the explanation mentioned above for the additive effect occurs in our mouse model of *S. aureus* bloodstream infection or other mechanisms are involved. Additionally, *S. aureus* also encodes for its arginase, which might as well behave like its host counterpart, thereby quenching away the L-arginine for NOS and eventually generating less amount of NO (16, 42, 43). The inhibitory effect of L-proline and branched-chain amino acids on staphylococcal arginase will need to be determined in future studies.

It is astounding that host employs L-proline and branched-chain amino acids as anti-infection metabolites against *S. aureus* bloodstream infection since *S. aureus* growth in media lacking L-proline, L-valine, or L-leucine shows an amino acid auxotrophy, albeit this bacteria genome contains entire gene sets for the biosynthesis of these amino acids (44, 45). The prevailing situation we can imagine upon staphylococcal infection is that the host should limit the production of these amino acids so that this pathogen is unable to have abundant nutrients to grow immoderately. However, the fact says differently, which exhibits the elevation of these amino acids in the serum of infected animals. Consistent with the observation, *S. aureus* infection reduces the transcriptional level of branched-chain amino acid transaminase 2 that mainly contributes to the degradation of branched-chain amino acids (46), suggesting the lower degradation rate of branched-chain amino acids in infected mice. Instead of aiding the growth of *S. aureus*, exogenous supplementation of L-proline and branched-chain amino acids facilitates the phagocytes-mediated OPK of *S. aureus* and the elimination of staphylococci in the host. The mechanism of how the host accumulates a high level of L-proline and branched-chain amino acids *in vivo* upon staphylococcal infection is unknown and will be determined in future studies. TLR2/TLR6 agonist stimulates the significant increase of L-valine and L-isoleucine in mouse serum sample (10), which provides the clue *that S. aureus*-derived lipoteichoic acid and peptidoglycan might play a role in the induction of L-proline and branched-chain amino acids in infected animals.

## Materials and methods

### Bacterial strains, culture conditions, and experimental animals

*S. aureus* strain Newman (ATCC 25904), USA300 (ATCC BAA-1717), or MRSA252 (ATCC BAA-1720) was cultured from frozen stocks in tryptic soy agar (TSA) at 37°C incubator. The single colony was grown in tryptic soy broth (TSB) in a shaker bath at 37°C. Overnight cultures were diluted 1:100 into fresh medium and harvested at an absorbance of 1.0 (OD_600_) by centrifugation at 6000 g for 10 min. The cells were washed and re-suspended in sterile PBS. Female BALB/c mice (6 weeks old) were reared in cages fed with sterile water and dry pellet diets. Each mouse was then intravenously infected by inoculation with the low (0.3 × 10^7^), moderate (0.7 × 10^7^), or high (1 × 10^7^) CFU of *S. aureus* Newman or with sterile PBS (no-infection group). Between 50 and 100 μL blood were collected from the orbital vein of each mouse at 12 h post-infection.

### Plasma metabolite extraction

The metabolite extraction procedure was performed following methods described previously (23, 47). In brief, 50 μL plasma was quenched by using 50 μL cold methanol and collected by centrifugation at 8,000 rpm for 3 min. This step was performed twice. The two supernatants were mixed, and an aliquot of sample was transferred to a GC sampling vial containing 5 μL 0.1 mg/mL ribitol (Sigma) as an internal analytical standard and then dried in a vacuum centrifuge concentrator before the subsequent derivatization. Two technical replicates were prepared for each sample. All animal experiments were performed following institutional guidelines following the experimental protocol review.

### Derivatization and GC-MS analysis

Sample derivatization and subsequent GC-MS analysis were carried out as described previously (22, 23). Briefly, 80 μL of methoxamine/pyridine hydrochloride (20 mg/mL) was introduced to dried samples to induce oximation for 1.5 h at 37°C, and then 80 μL of the derivatization reagent MSTFA (Sigma) was mixed and reacted with the sample for 0.5 h at 37°C. A 1 μL aliquot of the derivative of the supernatant was added to a tube and analyzed using GC-MS (Trace DSQ II, Thermo Scientific). For data processing, spectral deconvolution and calibration were performed using AMDIS and internal standards. A retention time (RT) correction was operated in all samples, and then the RT was used as a reference against which the remaining spectra were queried, and a file containing the abundance information for each metabolite in all samples was assembled. Metabolites from the GC-MS spectra were identified by searching in the National Institute of Standards, and Technology (NIST) library used the NIST MS search 2.0. The resulting data matrix was normalized using the concentrations of exogenous internal standards, which were subsequently removed so that the data could be used for modeling consisted of the extracted compound. The resulting normalized peak intensities formed a single matrix with Rt-m/z pairs for each file in the dataset. To reduce the between-sample variation, we centered the imputed metabolic measures for each tissue sample on its median value and scaled it by its interquartile range (48). ClustVis, a web tool for visualizing the clustering of multivariate data, was employed to create PCA plot and heatmaps (49). Metabolic pathways were enriched by utilizing MetaboAnalyst 4.0 (50).

### Effect of metabolites on *S. aureus* infection

Female BALB/c mice were acclimatized for three days and then randomly divided into groups for investigating the effects of L-proline, L-isoleucine, L-valine, L-leucine, or a mixture of four metabolites as mentioned above (L-proline, L-isoleucine, L-leucine, and L-valine, designated as PILV). Before the infection of *S. aureus* Newman, 100 μl of each metabolite (0.5 g kg^−1^) or an equal volume of sterile saline (no metabolite control) was intraperitoneally injected. 6h later, mice were intravenously challenged by *S. aureus* Newman (1 × 10^7^ CFU/mouse) and continued to be given the metabolites daily by intraperitoneal injection. Bodyweight of infected animals was measured daily. On day 15 following infection, mice were euthanized by CO_2_ inhalation and cervical dislocation. Both kidneys were separated, and bacterial load in one organ was detected by homogenizing tissue with PBS containing 0.1% Triton X-100. Serial dilutions of homogenate were sampled on TSA and incubated for colony formation overnight at 37°C. The remaining organ was investigated by histopathology analysis (51). For survival, *S. aureus* USA300 or MRSA252 were chosen. Six weeks old BLAB/c mice were intravenously inoculated with 100 μl of bacterial suspension in PBS at a concentration of 2 × 10^8^ CFU ml^−1^ (USA300) or 2 × 10^9^ CFU ml^−1^ (MRSA252). Each metabolite, PILV, BEC (S-(2-boronoethyl)-L-cysteine, arginase inhibitor, 50 mg kg^−1^), or both of nitric oxide (NO) inhibitor (L-NMMA or L-NAME, 40 mg kg^−1^) and PILV was given at the manner as mentioned earlier. PILV was administrated with 100 μl in PBS at a concentration of 0.5 g kg^−1^ for each metabolite. Survival was monitored over 14 days.

### Determination of NO release, urea and arginase activity

NO release in serum or tissues was calculated by examining the nitrate and nitrite concentrations with a Total Nitric Oxide Assay Kit (Beyotime, China) according to the manufacturer’s instructions. The optical densities at 540 nm were recorded using a Microplate Reader (Biotek Instruments, Inc., Vermont, USA). The concentration of NO output was calculated from the standard curve. Urea production in serum was determined using a Urea Colorimetric Assay Kit (BioVision). The mouse serum was collected for the Arginase Activity Assay kit (Sigma, MAK112).

### Quantitative real-time PCR

Total RNA was isolated from blood and tissues using TRIzol reagent, respectively (Invitrogen, Carlsbad, CA, USA). The cDNA was synthesized according to the manufacturer’s instruction of the PrimeScript^TM^ RT reagent Kit with the genomic DNA Eraser (Takara, Kyoto, Japan). Then, the mRNA levels of genes Arg1, Arg2, NOS1, NOS2, and NOS3 were detected using quantitative real-time polymerase chain reaction (qRT-PCR) with TB Green™ Premix Ex Taq™ II (Takara) in the LightCycler96 system (Roche, Indianapolis, IN, USA). The housekeeping gene β-Actin (ACTB) was used as an endogenous control. All primers are listed in Table S1. The qRT-PCR conditions were as follow 95 °C for 5 min followed by 45 cycles of 95 °C for 10 s, 58 °C for 30 s and 72 °C for 30 s. For the final melting curve step, the samples were subjected to 95 °C for 10 s and 65 °C for 1 min and then ramped to 97 °C by 5 °C every 1 s with a final cooling step at 37 °C. After three repeated PCRs, the gene expression levels were calculated using the 2^−△△^CT method (52).

### Bacterial survival in human and mouse blood

To measure bacterial replication and survival ex vivo, fresh human blood was collected with heparin, an anti-coagulated reagent. Prior to incubating with 50 μl of a live bacterial suspension containing 5 × 10^6^ CFU, 0.45 ml of human blood was pretreated by 50 μl of heat-killed *S. aureus* Newman (5 × 10^5^ CFU) at 37°C for 30 min. Then the human blood sample was mixed with live bacterial suspension in the presence or absence of PILV (10 mM for each metabolite), NO inhibitor (L-NMMA or L-NAME), or both. For mouse blood studies, 100 μl of heat-killed *S. aureus* Newman (5 × 10^5^ CFU) was intravenously injected into BALB/c mouse. 6 h later, whole blood was collected by cardiac puncture. 50 μl of a live bacterial suspension including 5 × 10^5^ CFU *S. aureus* Newman was mixed with 0.45 ml of mouse blood in the presence or absence of PILV (10 mM for each metabolite), NO inhibitor (L-NMMA or L-NAME), or both. All these samples were incubated at 37°C with slow rotation for 60 min. After that, 0.55 ml of lysis buffer (0.5% saponin, 200 U streptokinase, 100 µg trypsin, 2 µg DNase, 10 µg RNase per ml PBS) was added to each sample for 10 min at 37°C before plating on TSA for enumeration of CFU (53, 54).

### Cell culture and quantitative phagocytosis assay

The murine macrophage cell line RAW264.7 was cultured in DMEM supplemented with 10% (V/V) cosmic calf serum (HyClone), 100 U mL^−1^ penicillin G and 100 U/mL streptomycin. The human macrophage cell line U937 was grown in RPMI 1640 medium supplemented with 10% heat-activated fetal bovine serum. All cells were grown at 37°C in a 5% CO_2_ incubator. U937-derived macrophages were induced by 160 nM phorbol 12-myristate 13-acetate (PMA) at 37°C for 48 h. Macrophage phagocytosis was investigated as described previously (22, 23). Briefly, RAW264.7 cells were harvested using CaCl_2_- and MgCl_2_-free PBS containing 5 mM EDTA and plated at 5 × 10^6^ macrophages/well in 6-well plate. U937-derived macrophages were seeded at 1 × 10^6^ cells/well in a 12-well plate. For experiments with the administration of PILV, NO inhibitor, or both, the cells were deprived of serum overnight and then incubated with the molecules as mentioned above in serum-starved media. After pretreating for 6 h, FITC-conjugated *S. aureus* was centrifuged onto RAW264.7 or U937-derived macrophages at a multiplicity of infection (MOI) of 100 in the indicated medium without serum or antibiotics. Then, the plates were placed at 37 °C for 1 h. After incubation, the macrophages were vigorously washed with cold PBS to stop additional bacterial uptake or to destroy the bacteria in the phagosomes. Cells were washed at least four times in cold PBS and then fixed in 4% paraformaldehyde before being harvested in cold PBS containing 5 mM EDTA and subjected to FACS^®^ analysis.

### Statistical Analysis

The relative abundance of metabolites among different groups, staphylococcal survival in blood, NO and urea levels, or macrophage phagocytosis was analyzed with the two-tailed Student *t*-test. Bacterial loads and abscess numbers in renal tissues were analyzed with the two-tailed Mann–Whitney test. All data were analyzed by GraphPad Prism (GraphPad Software, Inc.), and *P* values < 0.05 were considered significant.

## Acknowledgments

This work was supported by the National Natural Science Foundation of China grant 31700119 and the Natural Science Foundation of Guangdong Province grant 2019A1515012211 to Jinan University, the GDIM Young Elite Talents Project grant GDIMYET20170104 to Guangdong Institute of Microbiology, the Shenzhen Science and Technology Innovation Commission Project grant JCYJ20170815113109175 and the Dapeng Project grant KY20170202 to Shenzhen International Institute for Biomedical Research.

## Author contributions

X.C., R.P., Y.S., developed methods and conceptualized ideas. R.P., Y.S., H.Z., X.C., designed experiments, X.C., R.P., Y.S., performed experiments, X.C., R.P., analyzed data, X.C., R.P., wrote the manuscript.

---

**Figure S1.**
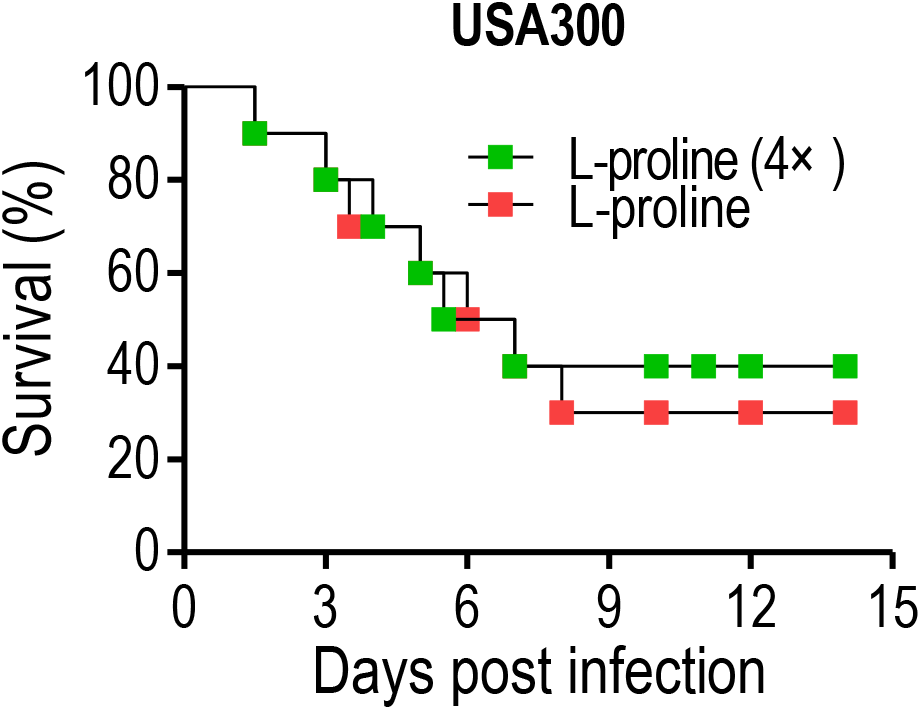
An increasing dose of L-proline is not able to increase survival.

**Figure S2.**
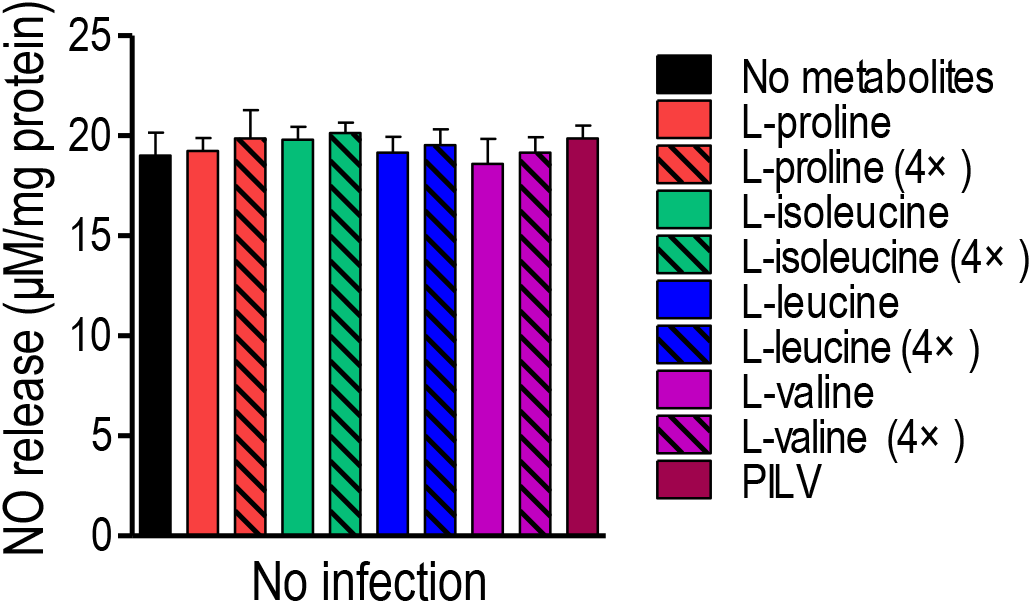
Metabolite treatments are not able to increase NO release in the mouse without *S. aureus* infection.

**Table S1.**
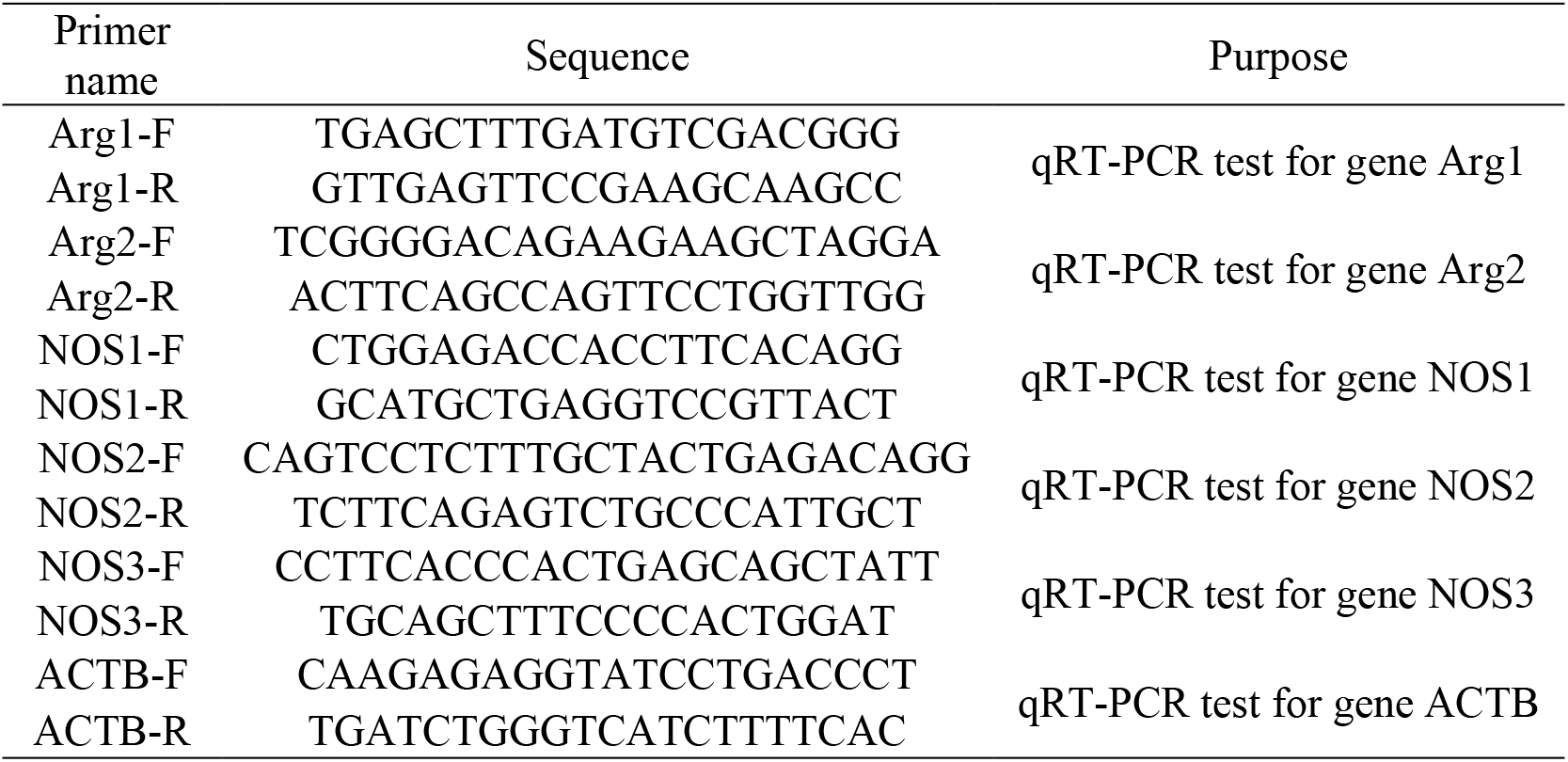
The primers used in this study

